## Supplementary Material for "Using genomic prediction to detect microevolutionary change of a quantitative trait"

### Electronic Supplementary Material

**Table S1**: Sample sizes in each year. Sample size refers to the number of genotyped individuals (training and test populations combined).

| Year | Sample Size |
|---|---|
| 1977 | 1 |
| 1978 | 1 |
| 1979 | 11 |
| 1980 | 9 |
| 1981 | 8 |
| 1982 | 10 |
| 1983 | 25 |
| 1984 | 26 |
| 1985 | 6 |
| 1986 | 34 |
| 1987 | 78 |
| 1988 | 33 |
| 1989 | 96 |
| 1990 | 127 |
| 1991 | 178 |
| 1992 | 176 |
| 1993 | 198 |
| 1994 | 186 |
| 1995 | 187 |
| 1996 | 208 |
| 1997 | 223 |
| 1998 | 246 |
| 1999 | 186 |
| 2000 | 218 |
| 2001 | 275 |
| 2002 | 147 |
| 2003 | 253 |
| 2004 | 303 |
| 2005 | 319 |
| 2006 | 230 |
| 2007 | 303 |
| 2008 | 282 |
| 2009 | 264 |
| 2010 | 303 |
| 2011 | 344 |
| 2012 | 317 |
| 2013 | 250 |
| 2014 | 328 |
| 2015 | 276 |

**Table S2: Summary of linear mixed models exploring phenotypic trends.** Models are shown for (a) 3622 measurements on 1259 animals born between 1976 and 2015 and (b) 1045 measurements on 401 animals born between 2005 and 2015 i.e. the cohorts born after the earlier study [1]. Note that birth year fitted as a random effect explains less than 3% of the variance in adult weight. Birth year fitted as a fixed effect reveals that adult weight significantly decreased over the entire study period, but the decline had ceased or even reversed since 2005. Numbers in parentheses are standard errors of model terms.

| | *Random Effects* | | | | *Fixed Effects* | | |
|---|---|---|---|---|---|---|---|
| Cohorts | Birth Year | Capture Year | ID | Residual | Sex | Age | Birth Year |
| (a) 1976-2015 | 0.31<br>(0.55) | 0.96<br>(0.98) | 6.13<br>(2.48) | 3.07<br>(1.75) | 9.51 (0.18)<br>$P < 2.x10^{-16}$ | 0.28 (0.02)<br>$P < 2.x10^{-16}$ | -0.067 (0.022)<br>P = 0.003 |
| (b) 2005-2015 | 0.15<br>(0.39) | 0.42<br>(0.65) | 4.67<br>(2.16) | 2.30<br>(1.52) | 9.65 (0.28)<br>$P < 2.x10^{-16}$ | 0.61 (0.07)<br>$P =1.4.x10^{-6}$ | 0.106 (0.085)<br>P = 0.23 |

**Table S3: Cohort mean genomic estimated breeding values regressed against birth year.** Cohort mean GEBV was regressed on year of birth, weighted by sample size in each year. We used both frequentist and Bayesian methods, with the former treating the posterior mean of each individual GEBV as a point estimate and the latter considering the uncertainty in GEBVs by performing regressions on the entire posterior distribution of GEBVs. The anti-conservative P values from the frequentist models are included to allow comparison with the previous study [1]. The Bayesian method, i.e. the method recommended by Hadfield and colleagues [2], is reported in the main text. $r^2$, F and P are from linear models using the posterior means of individual GEBVs to estimate cohort means. 95% CI is the credible interval for the coefficient of mean cohort GEBV regressed on year. $P_{MCMC}$ is the probability of the slope being less than zero, after accounting for the uncertainty in individual GEBVs i.e. by treating the posterior distribution of GEBVs rather than the posterior mean of GEBVs. $P_{drift}$ goes a step further by not only accounting for the uncertainty in individual GEBVs but also testing the probability that the trend is caused by genetic drift alone. This was performed by comparing observed slopes with slopes derived from gene-dropping simulations (see main text).

In the main text, the results shown are for the training and test populations in the cohorts born between 1990 and 2015 (italicised row). Restricting the analysis to only the test population would result in the same overall conclusions, as would considering the cohorts born between 1980 and 2015. Restricting the analyses to the 2005-2015 cohorts reduces the statistical support for an increase in GEBVs, because the estimated slopes of GEBV on year have a wider credible interval than for the longer time series.

| | | Frequentist method | | | | Bayesian method | | |
|---|---|---|---|---|---|---|---|---|
| Cohorts | Population | Coeff (SE) | $r^2$ | F | P | Coeff [95% CI] | $P_{MCMC}$ | $P_{drift}$ |
| 1980-2015 | Test<br>*n=5495* | 0.011<br>(0.002) | 0.46 | 30.30<br>(1,33) | **$4.16 \times 10^{-6}$** | 0.011<br>[0.0008-0.0207] | **0.014** | 0.059 |
| | Test + Training<br>*n=6652* | 0.010<br>(0.002) | 0.47 | 31.41<br>(1,34) | **$2.82 \times 10^{-6}$** | 0.010<br>[0.0015-0.0188] | **0.008** | 0.052 |
| 1990-2015 | Test<br>*n=5339* | 0.010<br>(0.002) | 0.44 | 20.86<br>(1,24) | **$1.25 \times 10^{-4}$** | 0.010<br>[0.0000-0.0208] | **0.024** | 0.068 |
| | *Test + Training*<br>*n=6327* | *0.011*<br>*(0.002)* | *0.50* | *25.60*<br>*(1,24)* | ***$3.57 x10^{-5}$*** | *0.011*<br>*[0.0009-0.0204]* | ***0.014*** | *0.057* |
| 2005-2015 | Test<br>*n=2819* | 0.010<br>(0.005) | 0.19 | 3.40<br>(1,9) | 0.099 | 0.010<br>[-0.0134-0.0309] | 0.184 | 0.243 |
| | Test + training<br>*n=3216* | 0.012<br>(0.006) | 0.24 | 4.21<br>(1,9) | 0.070 | 0.011<br>[-0.0102-0.0313] | 0.133 | 0.192 |

**Figure S1:** Histograms showing the distribution of the difference in slope of GEBV against year for the real data minus the equivalent slope from the gene-dropped data. The proportion of slope differences <0 gives the probability that the observed slope is caused by genetic drift (statistic $P_{drift}$ in Table S3). Fig S1D shows data for test and training populations from the cohorts born between 1990 and 2015 i.e. the same plot as Fig 1C in the main paper. Other combinations of cohorts (1980-2015, 1990-2015 and 2005-2015) and population (training and test population, or test population only) give broadly similar results (see also Table S3).

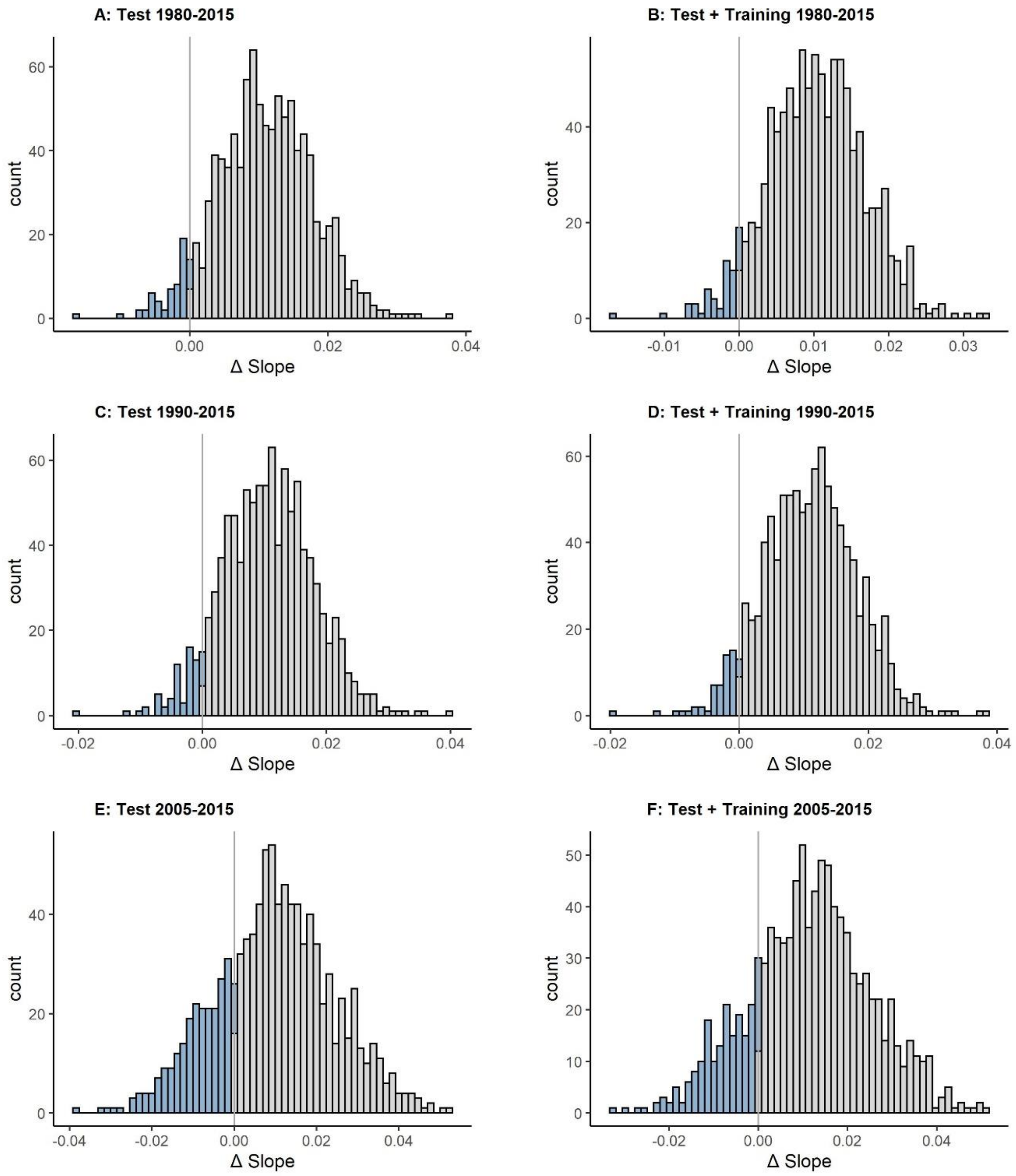

**Table S4: Using linear mixed models to test for microevolutionary change.** In the main paper we regressed cohort mean GEBVs against year of birth, the approach taken in most papers that have explored microevolutionary trends in breeding values. An alternative approach would be to use individual GEBVs and fit year of birth both as a fixed effect (to look for evolutionary trends) and as a random effect (to account for any between year heterogeneity in the variance of GEBVs). We ran linear mixed models, and the conclusions are essentially identical to those obtained from linear regressions of cohort mean GEBVs on year of birth. Note that birth year as a random effect explains less than 1% of the variance in GEBVs. The birth year as a fixed effect term gives very similar coefficients and statistical support as the regression of cohort mean GEBVs on year of birth (Table S3), using both frequentist and more conservative Bayesian methods.

| Cohorts | Population | Frequentist method | | | | | Bayesian method | | |
|---|---|---|---|---|---|---|---|---|---|
| | | Random effects | | Birth year as fixed effect | | | | | |
| | | Birth year | Residual | Coeff (SE) | df | P | Coeff [95% CI] | $P_{MCMC}$ | $P_{drift}$ |
| 1980-2015 | Test | 0.004 (0.060) | 0.625 (0.790) | 0.011 (0.002) | 28.9 | $5.63 \times 10^{-6}$ | 0.011 [0.0012-0.0208] | **0.006** | 0.051 |
| | Test + Training | 0.003 (0.057) | 0.686 (0.828) | 0.010 (0.002) | 32.7 | $2.93 \times 10^{-6}$ | 0.010 [0.0016-0.0183] | **0.006** | **0.049** |
| 1990-2015 | Test | 0.004 (0.063) | 0.625 (0.790) | 0.011 (0.002) | 21.9 | $9.87 \times 10^{-5}$ | 0.010 [0.0018-0.0211] | **0.020** | 0.065 |
| | Test + Training | 0.003 (0.056) | 0.678 (0.823) | 0.011 (0.002) | 22.4 | $3.20 \times 10^{-5}$ | 0.011 [0.0012-0.0205] | **0.010** | 0.055 |
| 2005-2015 | Test | 0.001 (0.027) | 0.602 (0.777) | 0.010 (0.005) | 8.45 | 0.098 | 0.010 [-0.0134-0.0311] | 0.179 | 0.183 |
| | Test + training | 0.001 (0.034) | 0.629 (0.793) | 0.011 (0.006) | 8.94 | 0.068 | 0.011 [-0.0101-0.0312] | 0.135 | 0.193 |

**Table S5: Regression of cohort mean GEBVs on year of birth using a 'leave one cohort out' genomic prediction analysis**. Genomic prediction models were run for each cohort, but with the focal cohort treated as the test population and all other cohorts as the training population. These GEBVs were then merged to create a dataset for all cohorts. This has the advantage that not only is the individual's phenotype not used to estimate its GEBV (thereby removing the risk that non-genetic effects on phenotype are biasing the GEBV), but neither are the phenotypes of any other animals from the same cohort. However, a major disadvantage is that the uncertainty in the GEBV estimates is harder to incorporate into the regressions of cohort mean GEBV on birth year. This is because the posterior distribution of SNP effects is different for each cohort. For this reason we cannot calculate $P_{MCMC}$ or $P_{drift}$. We do not report the P values for the regression of cohort mean GEBV on birth year because that would be anti-conservative due to the treatment of individual GEBVs as point estimates. Nonetheless, we note that the coefficients are slightly greater than in the equivalent models (training + test population) in Table S3. Thus, it is possible that the microevolutionary trends reported in the main text and Table S3 are slight underestimates.

| Cohorts | Population | Coeff (SE) |
|---|---|---|
| 1980-2015 | All (n = 6652) | 0.012 (0.002) |
| 1990-2015 | All (n = 6327) | 0.013 (0.002) |
| 2005-2015 | All (n = 3216) | 0.014 (0.006) |
